# Contingency awareness shapes neural responses in fear conditioning

**DOI:** 10.1101/2024.08.13.607803

**Authors:** Yuri G. Pavlov, Nick S. Menger, Andreas Keil, Boris Kotchoubey

## Abstract

Contingency awareness refers to an observer’s ability to identify the association between a conditioned (CS) and an unconditioned stimulus (US). A widely held belief in human fear conditioning is that this form of associative learning may occur independently of contingency awareness. To test this hypothesis, in this preregistered study (https://osf.io/vywq7), we recorded electroencephalography (EEG) during a task, where participants were presented with compounds of a word (drawn from two semantic categories) and tactile stimulation (vibration), followed by either a neutral sound (US-) or a loud noise (US+). Based on interviews, participants were divided into an aware (N=50) and an unaware (N=31) group. Only the aware group showed evidence of learning at the neural level, notably a larger stimulus-preceding negativity developing before US+ and a stronger theta response to vibrations predicting US+. The aware group also showed stronger alpha and beta suppression around the vibrations and a weaker theta response to US+, possibly indicating heightened attention to the cue and the violation/confirmation of expectation. Group differences in alpha and beta suppression were already present before the aversive learning began, suggesting that elevated attention may precede and facilitate awareness. Personality tests showed that elevated anxiety, neuroticism, higher intolerance of uncertainty, or harm avoidance are not predictive to the acquisition of contingency awareness. Our findings support the notion that fear conditioning, as reflected in cortical measures, cannot occur without contingency awareness.

## Introduction

The question of whether people can learn associations between events without consciously recognizing these relationships is a matter of debate (Labrenz et al., 2015). In the context of fear conditioning, the explicit knowledge of the conditioned-unconditioned stimulus (CS-US) association is referred to as contingency awareness. Despite decades of extensive research on conditioning, the importance of contingency awareness for fear learning remains debated. Some researchers argue that simple forms of associative learning are independent of awareness (Glenn et al., 2012; Jovanovic et al., 2006; Knight et al., 2003; Labrenz et al., 2015; Raio et al., 2012; Schultz & Helmstetter, 2010), while a multitude of evidence suggests that contingency awareness is a prerequisite for fear learning (Baeuchl et al., 2018; Glenn et al., 2012; Grillon, 2002; Klucken et al., 2009; Tabbert et al., 2011a). A comprehensive meta-analysis of thirty original studies found no compelling evidence supporting contingency-unaware fear conditioning (Mertens & Engelhard, 2019). Thus, two possibilities emerge: either fear memory can be formed without conscious recognition of the fear cues, or only consciously recognized threats can form long-term fear memories^1^. A deeper understanding of how individuals acquire contingency awareness and its impact on fear learning is vital for comprehending the psychopathology of fear.

Fear learning and the modulating effects of contingency awareness can be assessed through a variety of physiological measures. However, up to date adding these measures has not clarified the role of contingency awareness, because the results remain similarly inconclusive. For example, individual studies using skin conductance response (SCR) found that conditioned SCR can be produced in contingency-unaware participants (Bräscher & Witthöft, 2019; Cosand et al., 2008; Esteves et al., 1994; Knight et al., 2009), but in the majority of studies successful conditioning was observed only in participants who were aware of the contingencies (Baeuchl et al., 2018; Biferno & Dawson, 1977; Dawson et al., 1979, 2007; Grillon, 2002; Hamm & Vaitl, 1996; Klucken et al., 2009; Marinkovic et al., 1989; Merz et al., 2010; Purkis & Lipp, 2001; Sevenster et al., 2014; Tabbert et al., 2006). Studies of the neural correlates of contingency awareness also yield equivocal results. Again, most studies directly or indirectly indicate the involvement of awareness and attentional control in classical conditioning learning (Baeuchl et al., 2018; Carter et al., 2006; Lam et al., 2023; McIntosh et al., 2003; Tabbert et al., 2011a). On the other hand, unaware learning was shown to engage broad areas within the primary fear network of the human brain as well as sensory cortical areas (Klucken et al., 2009). Studies of metabolic responses reflected in fMRI BOLD signal are useful in increasing our understanding of the spatial distribution of activity in the brain related to contingency awareness, but the method has limited temporal resolution. Similarly, arousal measures extracted from SCR are limited in temporal resolution.

In contrast, electroencephalography (EEG) offers excellent temporal resolution. Many neurocognitive processes in fear conditioning unfold after stimulus onset and are missed by simple behavioral measures, yet EEG can reveal them through event related potentials (ERPs) and oscillatory activity. With EEG we can test whether changes in CS valence arise during early sensory processing, during later evaluative processing indexed by the Late Positive Potential (LPP), or during anticipation of the US indexed by the stimulus preceding negativity (SPN). ERPs provide time resolved neural conditioned responses that help separate stages of fear processing, but evidence on how contingency awareness shapes these signals is scarce. For example, an EEG study found no significant EEG difference in LPP in CS+/− contrast between aware and unaware subjects (Pastor et al., 2015). SPN is a slow component that begins about 200 to 500 ms before expected events and may index affective anticipation of the US, with more negative amplitudes reflecting stronger expectancy (Baas et al., 2002; Kausche & Schwabe, 2020; Pavlov et al., 2023; van Boxtel & Böcker, 2004). Expectancy ratings are commonly used to quantify learning in human fear conditioning (Constantinou et al., 2021; Gromer et al., 2025), and we hypothesize that SPN amplitude can potentially serve as a proxy for expectancy and learning, with a more negative SPN elicited before the US. In the frequency domain of EEG, suppression of alpha activity has been linked to directed attention to conditioned stimuli, with greater suppression in anticipation of the US (Bacigalupo & Luck, 2022; Farkas et al., 2024). Beyond alpha, source-localized anterior cingulate theta power is amplified during fear recall (Bierwirth et al., 2021; Sperl et al., 2019), linking frontal midline theta with the engagement of attentional networks. Additionally, the theta activity is shown to be related to the violation of expectation or level of surprise and can track the motivational value of the stimulus (Cavanagh et al., 2012). These neural markers of learning may provide a useful readout of the differences in conditioning between aware and unaware learning.

Personality traits related to emotional responsiveness, such as anxiety or neuroticism, may influence the likelihood of becoming aware of contingencies. Again, the empirical data remain controversial, with studies reporting a negative correlation between anxiety and awareness (at least as a trend with p = 0.06: (Grillon, 2002), a positive correlation (Rehbein et al., 2015) or a lack of relationship (Berg et al., 2022; Kindt & Soeter, 2014). Consequently, the effects of trait anxiety on contingency awareness require further investigation and validation. The low replicability of these findings may be related not only to limited sample sizes but also to a broader distribution of anxiety and neuroticism in unaware as compared with aware participants. Other personality traits potentially related to contingency awareness include a stronger tendency to find uncertainty distressing (low intolerance of uncertainty) and a stronger tendency to avoid harm, both of which prompt seeking cues to predict the source of harm, which is closely related to trait fearfulness (Panitz et al., 2018). All these possibilities require further investigation.

The primary limitation in studying the neural underpinnings of contingency awareness has been the inherent imbalance between participant groups. Contingency awareness rates vary widely across studies. For example, Li et al. (2025) reported variability from 47% to 97% in the proportion of participants who acquired the contingencies, depending on the experimental methodology. In high trial count designs, however, only a small fraction of participants typically remain unaware, for example 10% (Friedl & Keil, 2021) and 22% (Farkas et al., 2024). In addition, most previous studies that studied contingency awareness either used paradigms with a small number of trials that are not suitable for EEG, employed tasks that masked the CS, thereby altering its sensory properties, or added another task that might distort brain activity associated with awareness. To increase the number of contingency unaware participants in our study, we employed a paradigm developed in our pilot work (see Supplement) with compound CSs consisting of a combination of words (drawn from two semantic categories: animals or clothing) and vibrations (applied to a finger of either the left or right hand), followed by a conventional US (a loud noise). See Figure 1 for details.

**Figure 1.**
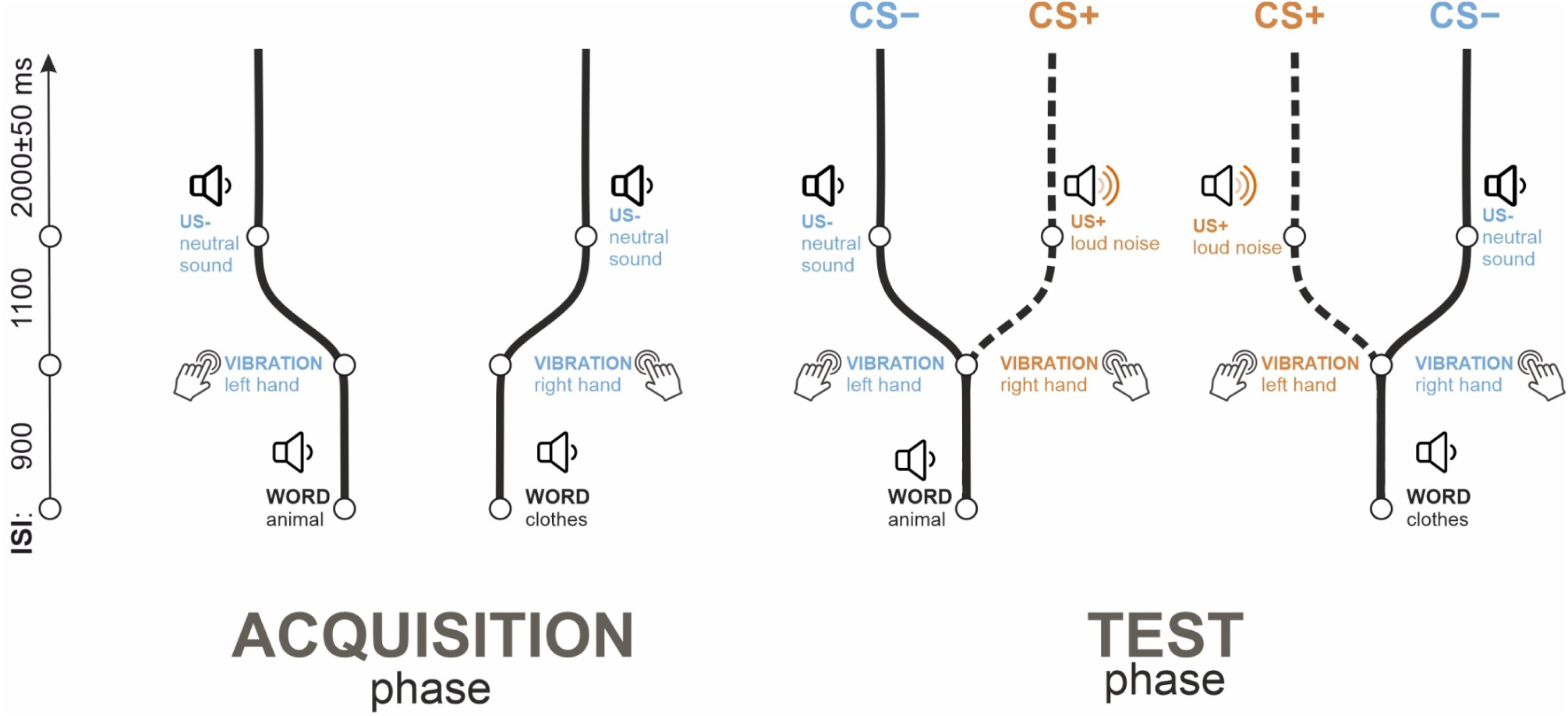
– Experimental design and individual trial structure. The experiment consisted of two phases: acquisition (left panel) and test (right panel). Each trial involved a sequence of three events: a word, followed by a tactile stimulus, and then an auditory stimulus. During acquisition, words from one category (e.g., animals) were followed by a left-hand vibration and a neutral sound (US-), while words from another category (e.g., clothing) were followed by a right-hand vibration and the same neutral sound. In the acquisition phase, there were 15 trials of each type. In the test phase, 110 trials replicated the acquisition sequences, and 110 trials presented incorrect word-tactile pairings followed by a loud noise (US+), designating the CS+ condition. Sequences with correct pairings were referred to as the CS-condition. During the acquisition phase only CS-sequences were delivered to decrease the complexity of the paradigm and facilitate learning while keeping the expected number of unaware participants high (as concluded from the pilot study; see Supplementary materials).

The aims of the current study were twofold:

(1) Leveraging the high temporal resolution of EEG signals, we aimed to explore the electrophysiological correlates of contingency awareness. To this end, we compared conditioned responses manifested in ERPs and oscillatory brain activity between groups of contingency aware and unaware participants. Specific hypotheses and the corresponding statistical tests were formulated based on a pilot study with 20 participants. A more detailed description of the pilot study is provided in the supplementary materials. In the current study, our first hypothesis (H1.1) stated that the difference between CS+ and CS-in SPN preceding the US (SPN_US_) would be greater in the aware group compared to the unaware group. Furthermore, SPN_US_ (H1.2) and SPN before the vibration (SPN_vibro_; H1.3) were expected to be generally larger in the aware group. Regarding time-frequency analysis and changes in the oscillatory brain activity, we had three hypotheses. The first related to alpha suppression preceding the second cue in the compound CS we used (word+vibration). We hypothesized stronger alpha suppression preceding the vibration (H1.5) in the aware group. Additionally, in response to tactile stimulation, there is typically strong suppression of alpha and beta activity over central and centro-parietal channels, often followed by a beta rebound. We observed these two events in the pilot sample data and formulated two hypotheses accordingly: we expected stronger beta oscillatory activity enhancement preceding the US (H1.4) and stronger alpha/beta suppression in response to the vibration (H1.6) in the aware group.
(2) We aimed to explore the role of individual differences in personality traits related to emotional processing in determining the probability of becoming aware of the contingencies. To this end, we compared the two groups in terms of their personality traits. These preregistered hypotheses were formulated in light of inconsistencies in the literature and our own reasoning and intuition, and they did not rely on pilot data. Specifically, we hypothesized that (H2.1) state anxiety would increase to a larger extent in the unaware group compared to the aware group in the pre-vs. post-acquisition comparison. Next, we hypothesized that anxiety scores would have broader distribution in the unaware group than in the aware group (i.e., more individuals with extreme anxiety/neuroticism scores would be in the unaware group, while most values in the aware group would be in the middle range). Thus, the distribution shape for anxiety (H2.2) and neuroticism (H2.3) would differ between the aware and the unaware group. Lastly, harm avoidance (H2.4) and intolerance of uncertainty (H2.5) were expected to be higher in the aware group.

## Materials and Methods

### Participants

Eighty-one participants provided complete datasets (56 females, 23 males, 2 others; mean ± SD, 28.5 ± 10.9 years). Based on structured interviews (described in detail below), participants were divided into two groups: aware (N=50) and unaware (N=31).

The study was advertised through mass emails at the University of Tübingen. The inclusion criteria included being between 18 and 59 years old and a native German speaker. The exclusion criteria were any past neurological or psychiatric diseases, taking any medication during the study. Participants gave their informed, written consent and were compensated with €10 or received course credit for their participation. The study was approved by the local Ethics Committee of the Faculty of Medicine of the University of Tübingen.

### Sample size justification

For the H1 hypotheses (Group and Condition effects on EEG variables), we aimed to achieve sufficient statistical power to detect an effect size that is 75% of the smallest Group effect obtained in the pilot study with the same design (the main effect of Group on SPN_US_ (aware vs unaware): 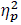 = .13, see below). Using G*Power, we conducted a power analysis for a one-way ANOVA with two levels of Group effect, 80% power, an alpha level of 0.05, and an effect size of 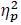 = 0.0975 (f = 0.328). A total of 76 participants, divided into two groups, would be sufficient to detect this effect.

For hypotheses H2.2-5 (group differences in personality traits) and H2.1 (group x pre-post interaction), we conducted a sensitivity analysis because we could not predict the effect size of these relationships. Similar to the analysis above, with 76 participants, we would be able to detect an effect size (main effect of Group or Group by PrePost interaction) of 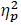 = 0.0975 (an equivalent of r = 0.31) with 80% power and alpha level of 0.05.

### Stimuli and procedure

#### Task

The experiment consisted of two phases: acquisition (Figure1, left panel) and test (Figure1, right panel). Each trial was a sequence of three events: presentation of a *word* followed by *tactile* stimulation followed by an *auditory* stimulus. The word+tactile stimulus compounds served as CS, and the auditory stimulus served as US.

During the acquisition phase, words from the first semantic category (e.g., animals) were consistently followed by a vibration applied to the left hand, and then by a neutral sound (US-). Similarly, words from the second semantic category (e.g., pieces of clothing) were paired with a vibration applied to the right hand, followed by the same neutral sound. The US presentation was followed by a silent interstimulus interval (ISI) lasting between 1950 and 2050 ms (2000±50 ms). The onset-to-onset interval between a word and a vibration stimulus was 900 ms, and that between the vibration stimulus and the US, 1100 ms. Each compound [semantic category 1 + left hand side] and [semantic category 2 + right hand side] was presented 15 times during the acquisition phase.

In the test phase, a total of 220 trials were presented. Of these, 110 trials replicated the sequences from the acquisition phase (55 starting with animal words and 55 with clothing words). In the other 110 trials, the semantic category + side rule was violated, meaning that categories were followed by a reversed tactile stimulus assignment (e.g., animal words followed by a right-hand vibration instead of a left-hand vibration). This reversed pairing was always followed by a loud noise (US+). Sequences containing the rule violation were designated as the CS+ condition, while the other sequences were referred to as the CS-condition.

Thus, in the test phase of the experiment, there were 110 trials in the CS+ condition (55 featuring left-hand side vibrations and 55 featuring right-hand side vibrations) and 110 trials in the CS-condition (also split into 55 left– and 55 right-hand side vibrations). These trials were included in the primary data analysis. The total duration of the experiment was approximately 17 minutes.

The order of the trials was pseudo-randomized, with the same sequence repeated no more than twice consecutively during the test phase. In the test phase, the same condition (CS+ or CS-) was not repeated more than four times consecutively. Each word appeared no more than 12 times throughout the experiment, with no repetitions on consecutive trials. The assignment of semantic categories to the side of the vibration was randomized between participants.

Participants were not given explicit instructions to learn the associations between the stimuli. They were simply instructed to remain seated with their eyes closed, attentively listen to the sounds, and perceive the vibrations. No behavioral responses were required during the task. However, the implicit task involved learning the association between the semantic categories, the side of the somatosensory stimulation, and the unconditioned stimulus (US). Throughout the experiment, participants were seated in a comfortable chair with armrests, placed in a soundproof, dimly lit EEG chamber.

#### Stimuli

The words were presented auditorily in German, with the duration of the word sounds varying between 413 and 772 milliseconds. These words were recorded using a female voice and were selected from two semantic categories: animals (Affe, Bär, Fisch, Gans, Hund, Katze, Kuh, Löwe, Pferd, Schwein, Schaf) and clothing (Anzug, Bluse, Handschuh, Hemd, Hose, Jacke, Jeans, Mantel, Rock, Socke, Strumpf). The average loudness of the words was 60 dB SPL (sound pressure level).

Tactile stimulation was delivered by two vibration units (10×3 mm vibromotors, typically used in cellphones), controlled via an Arduino Uno – a programmable open-source microcontroller board (D’Ausilio, 2012). These vibration units were attached to the distal phalanx of the middle fingers of both hands. The duration of the vibrations was 300 ms.

The aversive stimulus (US+) was a 1000 Hz sine wave embedded into a burst of white noise (92 dB, 500 ms). The US-was a neutral sound consisting of mixed frequencies (70 dB, 500 ms). The stimuli are available on OSF (https://osf.io/xph69/).

Before the experiment, participants familiarized themselves with the loud noise stimulus by listening to it three times with a 1-second SOA. Following the noise presentation, they were asked to rate the stimulus using Self-Assessment Manikins (SAM) on valence and arousal scales. These SAM ratings for the US+ were collected again after the experiment, subsequent to the contingency awareness interview. No SAM ratings were collected for CSs.

All auditory stimuli were presented through pneumatic earphones (3M E-A-RTONE).

#### Definition of awareness

Contingency awareness was assessed with a structured, hierarchical post experiment interview following the approach of Weike, Schupp, and Hamm (2007). The questions were adapted from Mann et al. (2001). The interview moved from open ended to specific probes to minimize demand characteristics and to avoid inducing awareness during assessment.

1. Participants were asked to describe what they heard and felt (e.g., words, sounds, left and right vibrations),
2. whether any words were predictive of the upcoming loud noise,
3. whether a vibration to the left or right hand predicted the loud noise,
4. whether a specific combination of words and vibration predicted the loud noise.
5. Participants were shown a schematic representing the experiment’s structure and asked if they recognized the connections between the words, vibrations, and sounds.

Typically, prompting participants to describe their experience in the first question, if they were contingency aware, resulted in a full report describing the experimental design (i.e., one semantic category was associated with a vibration applied to one hand, and a different category was associated with the other hand, where a violation of this association triggered the loud noise). If participants did not report the relationship between the stimuli after the first question, the next question was posed, and so on. Participants were considered “aware” if they could accurately describe the experimental design by the fourth question at the latest. The fifth question served for debriefing purposes. Additionally, participants who were identified as “aware” were asked at what point during the experiment they became aware of the contingencies.

The post-experiment interviews were consistent with a dichotomous pattern of contingency awareness. Most aware participants stated the CS-US relation in response to the first open-ended question, and a small number did so only after the second probe. Unaware participants reported no relation and typically expressed surprise when shown the schematic during debriefing.

#### Questionnaires

We employed four questionnaires to elucidate the relationship between probability to become aware of the contingencies and personality traits: (1) harm avoidance scale from the tridimensional personality questionnaire (Cloninger, 1987); (2) NEO-Five-Factor-Inventory (NEO-FFI) (Kanning, 2009); (3) intolerance of uncertainty (IUS) scale (Freeston et al., 1994); (4) State-Trait Anxiety Inventory (STAI) (Spielberger, 1970). The STAI-S (state anxiety) portion of the questionnaire was administered before and after conditioning. Remaining questionnaires were administered before conditioning. Overall scores on harm avoidance, IUS, neuroticism from NEO-FFI, and STAI entered the analyses.

#### Electroencephalography

A 64-channel EEG system with active electrodes (ActiCHamp, Brain Products) was used for the recording. The electrodes were positioned according to the extended 10-20 system, with Cz channel serving as the online reference and Fpz as the ground electrode. Impedance levels were kept below 25 kOm. The sampling rate was set at 1000 Hz.

Data preprocessing was carried out using EEGLAB (Delorme & Makeig, 2004). Recordings were filtered with a 0.1 Hz high-pass filter and a 45 Hz low-pass filter (using pop_eegfiltnew function with default parameters). Independent Component Analysis (ICA) was then performed using the AMICA algorithm (Palmer et al., 2012). As a preliminary step for improving ICA decomposition, a high-pass filter of 1 Hz was applied to the raw data (without a low-pass filter). The ICA results from the 1 Hz filtered data were subsequently applied to the data filtered with 0.1 Hz high-pass and 45 Hz low-pass filters. Components related to eye movements and high-amplitude muscle activity were removed, as were components mapped to a single electrode that could be clearly distinguished from EEG signals. The data were segmented into epochs from –1500 to 4000 ms, with 0 corresponding to word onset. Epochs containing remaining artifacts were visually identified and discarded.

For ERP analyses, baseline correction was applied to the data within the –200 to 0 ms interval. Before statistical analysis, the data were re-referenced to the average of the mastoids.

For the time-frequency analysis, all epochs were converted to current source density (CSD) using the CSD toolbox (Kayser, 2009). The spherical spline surface Laplacian was used with the following parameters: 50 iterations, m = 4, and a smoothing constant λ = 10^−5^ (Tenke & Kayser, 2005). Time-frequency analysis was conducted using the Fieldtrip toolbox (Oostenveld et al., 2011). Preprocessed single-trial data between 1 and 45 Hz in 1 Hz steps were transformed into the time-frequency domain using Morlet wavelets, with the number of cycles varying from 3 to 12 across 45 logarithmically spaced steps. Individual trial representations were then averaged across trials separately for each participant and condition. The analysis time window was shifted in steps of 20 ms. Spectral power in the time-frequency analysis was baseline-normalized by calculating the percent change relative to the –400 to –100 ms interval.

#### Statistics

We explored how ERPs, more specifically stimulus-preceding negativity (SPN), and changes in spectral power in theta, alpha, and beta frequency bands at different time points (i.e., before presentation of the US, preceding the vibrations, or in response to the vibrations) differed between conditions (CS+ vs CS-) and groups (aware vs unaware). The exact definitions of frequency bands, channels, and time windows are given in Table 1. Each region of interest (ROI) for averaging were specified a-priori based on the pilot study results.

**Table 1.** – Specification of ROIs.

| Hypothesis | Parameter | Channels | Frequency, Hz | Latency, ms |
| --- | --- | --- | --- | --- |
| H1.1, H1.2 | SPN preceding US | FC1, FCz, FC2, C2, C1, Cz | - | 1500-2000 |
| H1.3 | SPN preceding vibration | FC1, FCz, FC2, C2, C1, Cz | - | 500-900 |
| H1.4 | Beta preceding US | FC3, FC1, FCz, FC2, FC4, C3, C1, Cz, C2, C4, F3, F1, Fz, F2, F4 | 15-22 | 1500-2000 |
| H1.5 | Alpha preceding vibration | Pz, CPz, CP1, CP2, CP3, CP4, P1, P2, P3, P4 | 8-14 | 500-900 |
| H1.6 | Alpha/beta to vibration | Left hemisphere: C3, C1, CP1, CP3,<br>Right hemisphere: CP2, CP4, C2, C4 | 8-22 | 1000-1400 |
| Exploratory | Theta to vibration | Fz | 3-7 | 1000-1400 |
| Exploratory | Theta to US | Fz | 3-7 | 2000-2400 |

For testing H1 and H2.2, H2.4, H2.5, we used mixed ANOVAs with factors Condition (CS+/CS-) x Group (aware/unaware) and neural measures (see parameters in Table 1) as dependent variables.

Concerning hypotheses related to the personality traits and awareness associations, for testing H2.1, we used a mixed ANOVA with factors PrePost (before/after the conditioning) and Group (aware/unaware), with state anxiety as the dependent variable. For testing H2.3 and H2.2, we compared distributions of anxiety and neuroticism between the aware and unaware groups with Kolmogorov-Smirnov tests for the shape of the distribution and the Levene test for directed comparisons of variance in the two groups. Mean intolerance of uncertainty (H2.5) and harm avoidance (H2.4) values were compared between groups using one-way ANOVA.

We used p < .05 to determine statistical significance.

## Results

### Self-report data

Arousal (mean±SD, 7.34±1.69 on 1-9 scale) and valence (7.45±1.39 on 1-9 scale where 9 is the most unpleasant) ratings of the US+ (loud noise) did not differ between the groups (main effect of Group; arousal F(1, 79) < 0.01, p = .997, 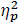 < .01; valence: F(1, 79) = 2.63, p = .109, 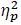 = .03), and no significant interactions were found. The US+ was rated as significantly more unpleasant after conditioning than before (main effect of Time (pre-post): F(1, 79) = 6.68, p = .012, 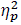 = .08; pre: 7.22±1.39, post: 7.68±1.37), probably indicating a sensitization effect.

### Event-related potentials

First, we assessed whether the stimulus preceding negativity (SPN) developed in anticipation of the vibrations (H1.3) would differentiate aware and unaware participants. As shown in Figure 2a, shortly after the presentation of the words, the SPN became larger in the aware group compared to the unaware group (see Table 2 for the full statistical output).

**Figure 2.**
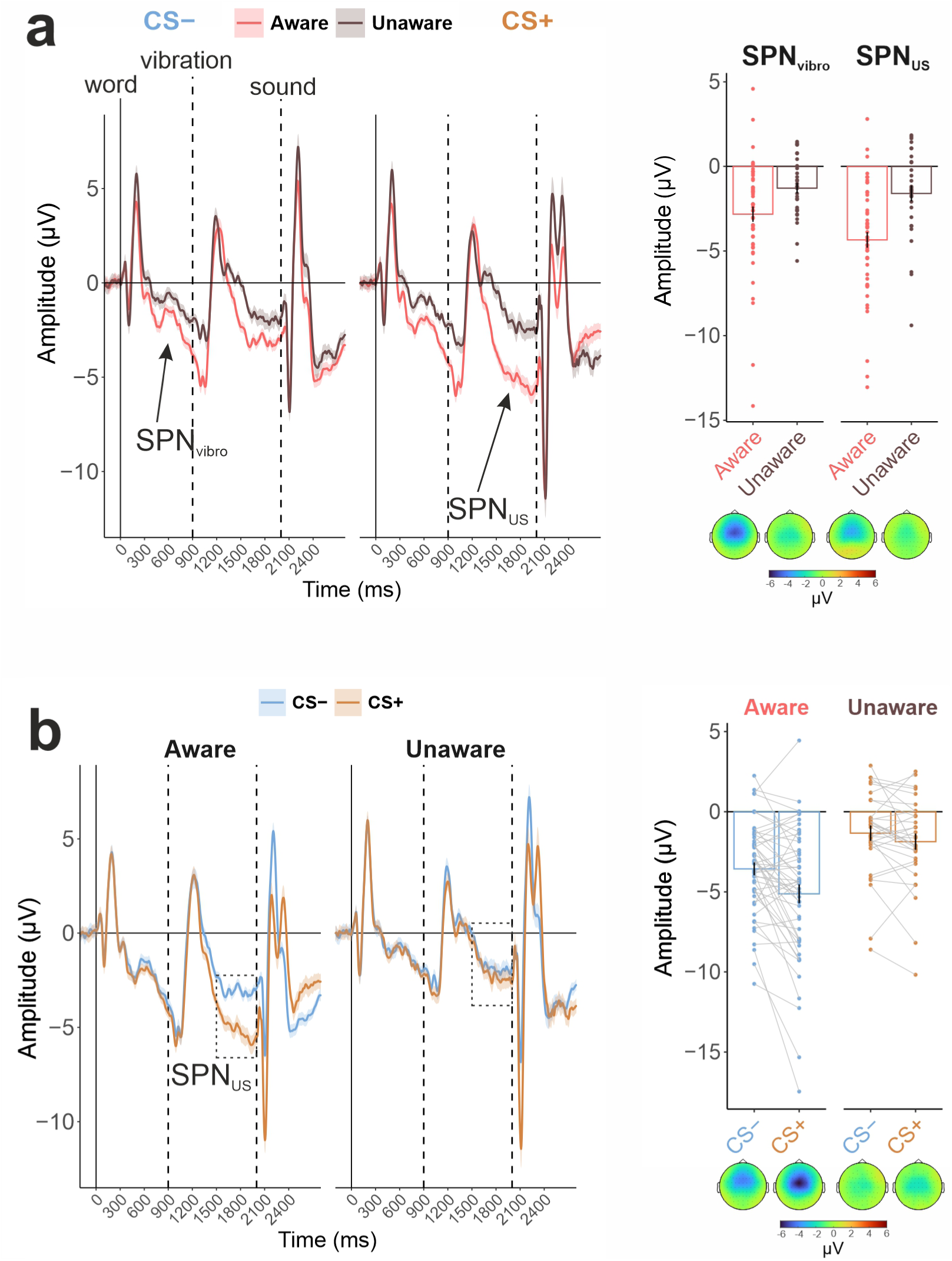
– The results of the event-related potentials analysis. All ERP curves are shown at Cz channel. (a) Comparison of the conditions in separate groups. The topographical maps show distribution of the amplitude averaged in the SPN_US_ time window (1500-2000 ms after word presentation). (b) Comparison of the groups in separate conditions. Shading is the Standard Error of the Mean (SEM).

**Table 2.**
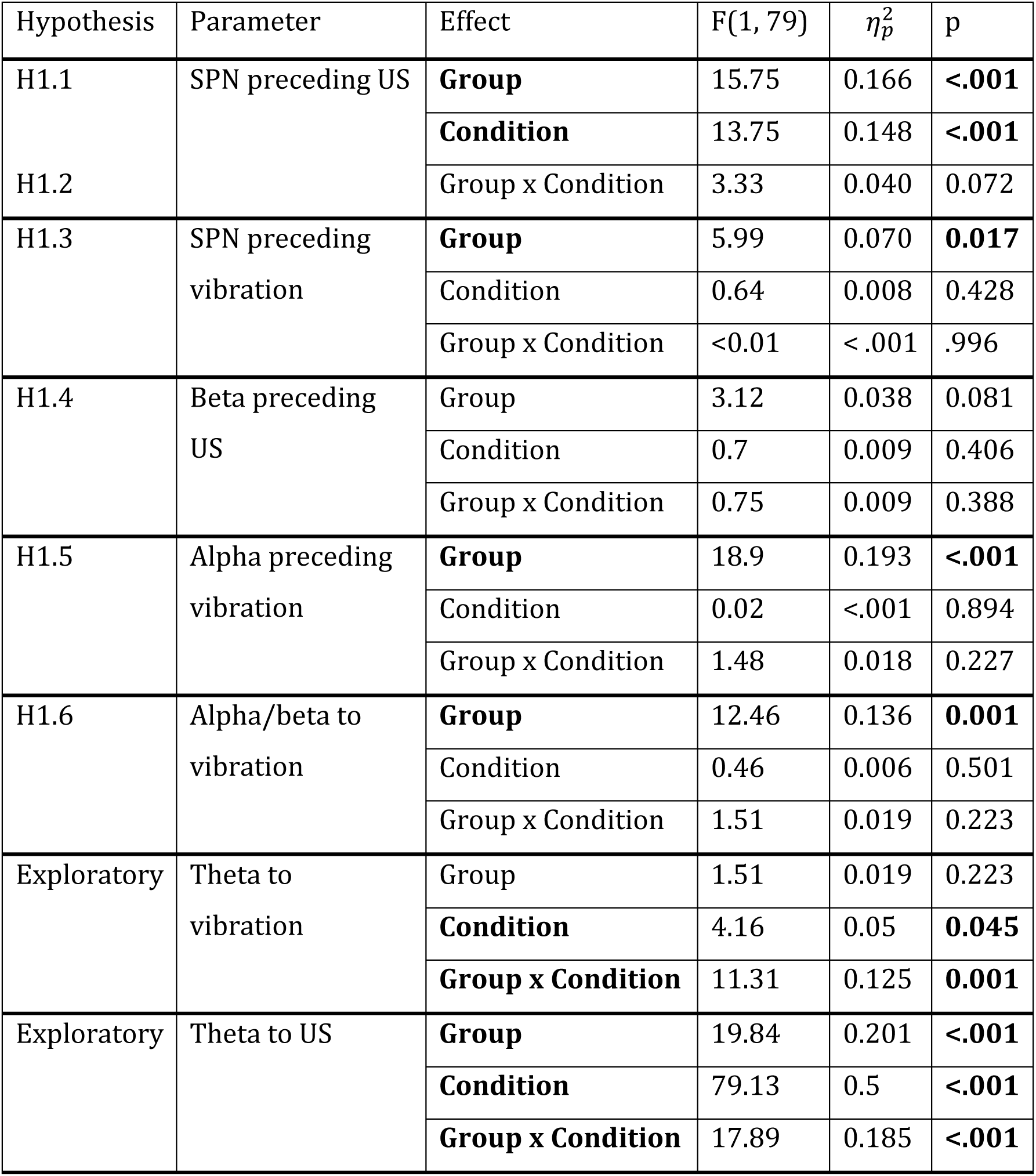
– The results of ANOVA.

Second, the SPN preceding the US was also larger in the aware group (H1.1). Additionally, the difference between the SPN preceding the loud noise (US+) and the SPN preceding the neutral sound (US-) was only present in the aware group (H1.2). The effect of condition was significant in the aware group (t(49) = 3.93, p < 0.001, d = 0.56) but not in the unaware group (t(30) = 1.67, p = 0.105, d = 0.30). However, this interaction effect (Group x Condition) was not significant in the preregistered cluster of electrodes. This discrepancy between the visual representation of the effect of condition in the groups and the statistical tests can be explained by the fact that the SPN preceding the US was primarily localized in fronto-central channels, while our preregistered cluster also included frontal channels. The effect of condition had a more central spatial distribution, with the interaction effect being significant only at Cz, FCz, C2, and FC2 channels.

A supplementary analysis of sensory ERPs to US (quantified as the N1-P2 amplitude difference to account for the baseline drift due to SPN) revealed a significant difference in sensory responsiveness between the groups with larger N1-P2 complex amplitude in the unaware participants (F(1, 79) = 8.81, p = 0.004, 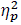 = 0.10). As expected, the amplitude in response to the loud noise (US+) was larger compared to the control sound (US-) (main effect of Condition: F(1, 79) = 11.74, p < 0.001, 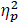 = 0.13). However, the interaction Group x Condition was not significant (F(1, 79) = 2.31, p = 0.133, 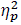 = 0.03).

### Time-frequency analysis

As shown in Table 2 and Figure 3, alpha activity preceding the vibrations and combined alpha and beta frequency band responses to the vibrations decreased stronger in the aware group compared to the unaware group. The beta rebound effect after the vibrations did not differ between the groups. Thus, the results supported hypotheses H1.5 and H1.6, but not H1.4.

**Figure 3.**
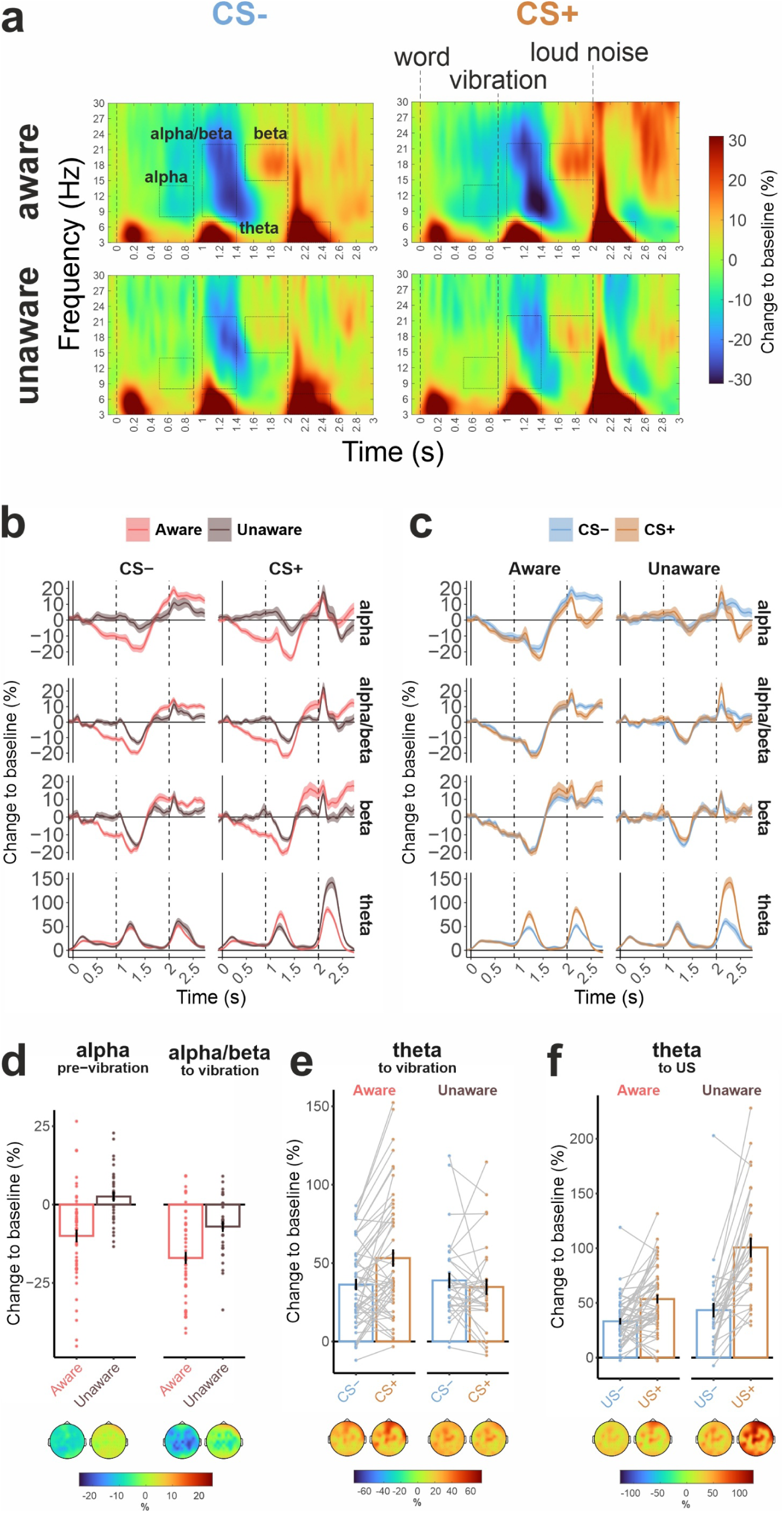
– The results of the time-frequency analysis. (a) Time-frequency maps with baseline-normalized spectral power averaged at all used channels (CPz, CP1, CP2, CP3, CP4, Pz, P1, P2, P3, P4, FC3, FC1, FCz, FC2, FC4, C3, C1, Cz, C2, C4, F3, F1, Fz, F2, F4). ROIs corresponding to tested hypotheses are highlighted with boxes. (b, c) Temporal dynamics of averaged in ROIs reported in Table 1. The vertical bars at 0, 0.9, and 2 s are corresponding to the word, vibration, and sound onsets. Shading is the SEM. (d, e, f) Mean baseline normalized spectral power value distribution for significant effects and the topographical distribution of the values over all channels.

Next, in a set of exploratory analyses, we tested the effects of condition and group on frontal midline theta. We found that the increase in theta power was stronger in response to the vibrations predicting the presentation of the loud noise (CS+ condition) compared to CS-, but only in the aware group (Condition x Group interaction; effect of Condition in Aware: t(49) = 4.39, p < 0.001, d = 0.63; Unaware: t(30) = 0.83, p = 0.412, d = 0.15; see Table 2 and Figure 3c and 3e).

Further differences between the groups were found in response to the US itself. While the main effect of the type of US was generally strong in both groups, the theta response to the loud noise was significantly stronger in the unaware group than in the aware group, whereas the response to the neutral sound was comparable between the groups (Group x Condition interaction; effect of Group for CS-: t(79) = 1.58, p = 0.119, d = 0.36; CS+: t(30) = 5.29, p < 0.001, d = 1.19).

To better understand what the primary factor was influencing group effects – contingency awareness or the inherent individual differences in stimulus processing that existed prior to the test phase and aversive conditioning – we examined the impact of group membership during the acquisition phase of the experiment (which occurred before any aversive stimulus was presented). This analysis revealed a stronger suppression of alpha (F(1, 79) = 23.99, p < 0.001, 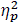 = 0.23) before the vibration, alpha/beta (F(1, 79) = 4.59, p = 0.035, 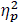 = 0.05) in response to the vibration, and beta (F(1, 79) = 4.01, p = 0.049, 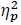 = 0.05) after the vibration in the aware group (see Supplementary Figure S2). However, no significant group effect was observed in the theta frequency band in response to the vibrations (F(1, 79) = 2.21, p = 0.141, 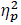 = 0.03).

### Personality measures

We hypothesized that trait anxiety and neuroticism would differ between the groups in terms of value distribution, with a wider distribution expected in the unaware group (i.e., more participants with either very high or very low levels of anxiety). However, comparisons of within-group variance using either Levene’s or Kolmogorov-Smirnov tests did not reveal significant differences (see Figure 4). Additionally, the mean values for intolerance of uncertainty and harm avoidance were not different between the groups (Table 3).

**Figure 4.**
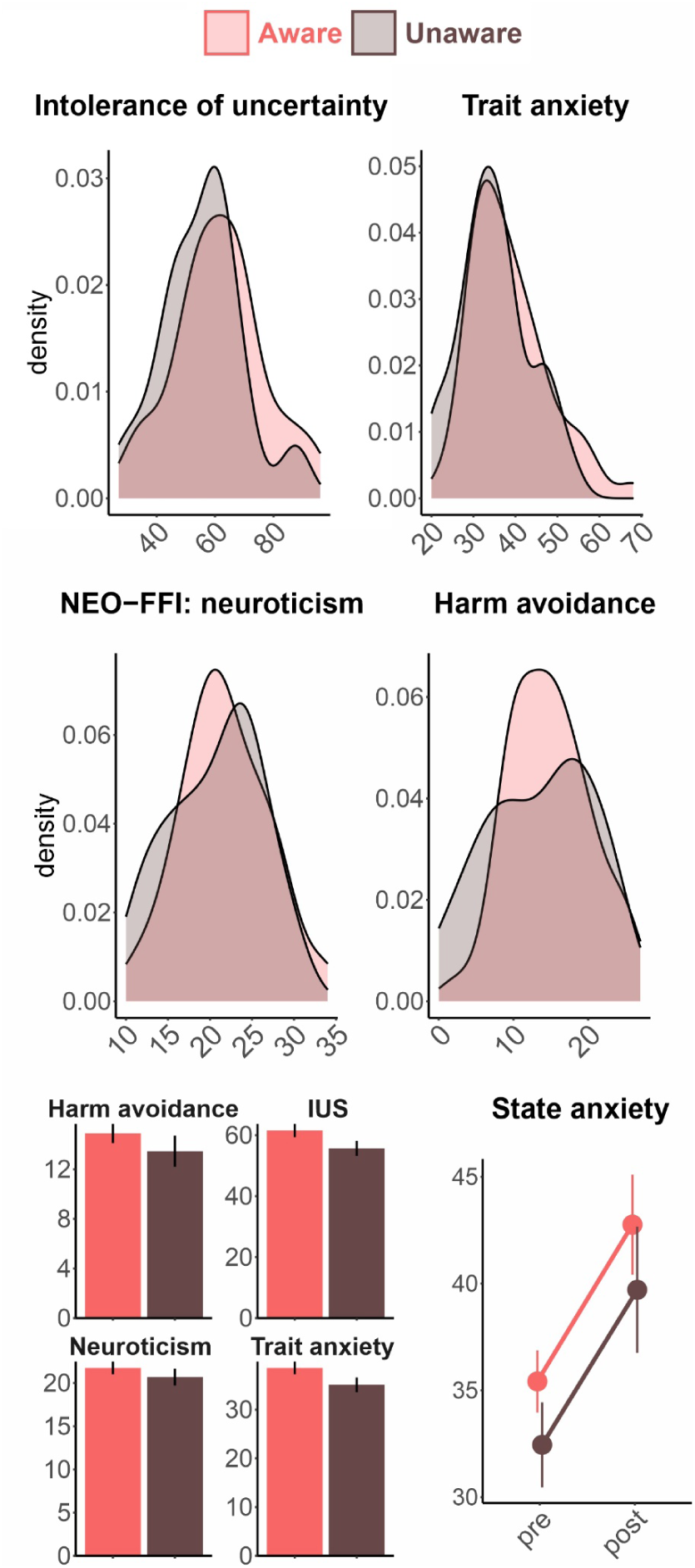
– Distribution of personality trait values in the aware and unaware groups.

**Table 3.** – The effect of group on personality traits.

| | $F(1, 79)$ | $\eta_p^2$ | $p$ |
| --- | --- | --- | --- |
| STAI trait | 2.847 | 0.035 | 0.096 |
| Harm avoidance | 1.05 | 0.013 | 0.309 |
| Intolerance of uncertainty | 2.996 | 0.037 | 0.087 |
| Neuroticism | 0.768 | 0.010 | 0.384 |

State anxiety increased after conditioning in similar amount in both groups (main effect of Time (pre-post): F(1, 79) = 44.45, p < 0.001, 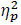 = 0.36; Time x Group interaction: F < 0.01; see Figure 4).

## Discussion

Using a novel fear conditioning paradigm, we tested the hypothesis that contingency awareness is necessary for fear learning in humans and explored multiple electrophysiological correlates of the psychological processes involved. The primary characteristic of awareness was found to be an enhancement in stimulus processing, as evidenced by stronger induced oscillatory responses and more negative stimulus-preceding negativity, indicating heightened anticipation of both CS and US. These anticipatory effects were further supported by attenuated theta responses to the US+ (loud noise) in the aware participants. Additionally, we examined whether individual differences in personality traits influence the likelihood of becoming aware of the contingencies. Our hypotheses regarding a wider distribution of anxiety and neuroticism or diminished intolerance of uncertainty and harm avoidance in unaware participants were not confirmed, suggesting that these personality traits have no or only minimal influence on the modulation of contingency awareness in fear learning.

We did not find evidence supporting the ability to learn associations between conditioned and unconditioned stimuli without contingency awareness. This finding corroborates a number of previous studies employing fMRI and peripheral measures such as SCR (Baeuchl et al., 2018; Klucken et al., 2009; Lovibond & Shanks, 2002; Tabbert et al., 2006), and aligns with evidence from behavioral studies (Mertens & Engelhard, 2019). The difference in SPN, an index of anticipation, indicated that only aware participants varied their expectancy of threat and safety. SPN is often largest at electrode Cz before the presentation of an expected auditory or somatosensory stimulus (Brown et al., 2008), which is where we found the interaction effect to be significant. However, the interaction effect did not reach the .05 level of significance in our study’s preregistered cluster of electrodes.

While SPN showed only weak sensitivity to differences in learning (CS+/CS-contrast) between aware and unaware participants, the theta oscillatory response to the vibrations demonstrated a strong interaction effect. The larger theta power in response to CS+ compared to CS-may be attributed to the violation of expectations that formed only in the aware group. When individuals are consciously aware of the link between word semantics and the side of the vibration, then receiving the vibration on the “wrong” side elicited a violation of expectations and associated frontal midline theta activation. Within some previous studies, a similar induced theta activity was observed in the tasks eliciting feedback-related negativity (FRN; Miltner et al. 1997; Cavanagh et al. 2011). Stronger frontal midline theta was also shown in case of aware recognition of the error (Wang et al., 2020), a signal coming from the same family of error responses. Furthermore, frontal midline theta is frequently interpreted as a signal of negative prediction error (Hajcak et al., 2006) or expectation violation (Cavanagh et al., 2011) because it is larger after punishment than after reward. Both FRN and the related theta responses are associated with sources in the anterior cingulate cortex (ACC) and the medial prefrontal cortex (mPFC; Miltner et al. 1997; Holroyd et al. 1998; Dehaene et al. 2003; Trujillo and Allen 2007; Beldzik et al. 2022). Moreover, the ACC is part of a distributed conscious control network that works alongside the prefrontal cortices in regulating conscious self-control – research indicates that a big portion of the activation of the ACC related to conflict happens only when conflicting stimuli are consciously perceived (Dehaene et al., 2003).

The stronger theta-band response to the US in the unaware group may indicate that theta activity is associated with expectation violation. In the aware group, the theta response to the loud noise was significantly reduced compared to the unaware group. This suggests that aware participants were prepared to handle unpleasant stimuli, while unaware participants experienced a consistent element of surprise. The lower theta response in the aware group may be due to their anticipation of the unpleasant US following the CS+, resulting in a relatively low prediction error. This could explain the reduced theta response to the loud noise. Alternatively, the weaker theta response to the US in aware participants might reflect a shift of theta activity from the US to the CS+ as part of the conditioning process. Although this interpretation cannot be ruled out, it would suggest that the CS+ acquires the aversive properties of the US, making the US less unpleasant for participants who exhibit a learning effect. However, in our study, aversiveness ratings for the US were identical in both aware and unaware individuals. Additionally, the increase in aversiveness ratings from the beginning to the end of the experiment was the same for both groups.

Expected sensory events are typically associated with lower neural sensory signals (Blakemore et al., 1998; Kok et al., 2012). A diminished sensory cortical response to painful stimulation when it is expected has been observed in previous EEG studies (Iannetti et al., 2008). The ERPs and, likely, the theta response do not capture the unpleasantness of the unconditioned stimulus per se but rather its saliency and capacity to attract attention (Iannetti et al., 2008). Meanwhile, the ability to anticipate the noxious stimulus did not make the loud noise more unpleasant – behavioral ratings of the US did not differentiate between the groups. Thus, noxious stimuli that is anticipated do not lose their potency to cause aversive feelings overall, but it does generate weaker cortical responses.

Previous fMRI studies showed an involvement of cingulate cortex (Lam et al., 2023) and insula (Bäuchl et al., 2018; Tabbert et al., 2011b) in aware conditioning. These brain areas are considered substrates for frontal midline theta activity (Asada et al., 1999; Ishii et al., 1999) and SPN (Hackley et al., 2020; Kotani et al., 2015), respectively, that we observed in the current study. Moreover, generally, anticipation of pain is tightly linked to BOLD activation in insula and cingulate cortex (Atlas, 2023; Palermo et al., 2015), as well as during anticipation of other kinds of aversive events (Andrzejewski et al., 2019). We can hypothesize that, similar to pain anticipation, insula – SPN, and cingulate cortex – theta are activated in conscious anticipation of an aversive loud noise.

In addition to the awareness-specific differentiation learning effects discussed above (and manifested in Condition x Group interactions), some other effects characterized general responsivity of aware participants (and were manifested in main Group effects). Specifically, aware individuals showed more negative SPN before both the US and the second stimulus in the CS compound (vibration). This suggests awareness-related overall enhanced anticipation of all relevant for learning stimuli. Similarly, in the context of non-aversive learning, it has been shown that CNV (which shares sources with SPN) is larger in participants who learned the S1-S2 association in a paradigm with masked S1 (Rozier et al., 2020).

The aware and unaware groups also exhibited different oscillatory responses to stimuli, with stronger alpha and beta suppression around the vibration, both before and after it occurred, in the aware group. The alpha suppression preceding the vibration likely reflects heightened attention in anticipation of the vibration, due to the pivotal role of the vibration side in determining the trial’s outcome. In contrast, the unaware participants did not show any attention enhancement preceding the vibrations with alpha activity returning to baseline level right after presentation of the first cues in the CS compound (words). We believe that alpha suppression can be seen as a general indicator of readiness to process the stimulus (Hoxha et al., 2023). Similarly, the stronger somatosensory response to tactile stimulation in the aware group may have facilitated more in-depth processing of the stimulus. Supporting our findings, aversive anticipation, compared to neutral anticipation, has been shown to engage larger areas in the visual and motor cortex (Andrzejewski et al., 2019), regardless of the stimulus modality.

The strong main effects of group (aware vs. unaware) on SPN, as well as on alpha and beta oscillations, suggest a significant involvement of attentional and sensory processes in contingency awareness. However, the direction of these relationships remains unclear. It is possible that contingency awareness amplifies the attentional processing of relevant stimuli. Conversely, the opposite pattern is equally possible: a high level of attention may underlie the ability to notice the CS-US contingency. The stronger suppression of alpha and beta activity in the acquisition phase of the experiment, which indicates enhanced attention to the stimuli even before the aversive learning began in the aware group, supports the latter explanation.

None of our hypotheses linking personality traits to the acquisition of contingency awareness were confirmed. Although individuals more affected by uncertainty might be expected to seek out contingencies more actively, this was not supported by the analysis of individual differences in intolerance of uncertainty. The same was true for trait anxiety, harm avoidance, and neuroticism. It is likely that other situational factors and traits related to cognitive functioning, such as curiosity and a tendency to mind-wander, are more predictive of the acquisition of contingency awareness. Future studies will provide further insights.

### Limitations

The present work is not free from limitations. Together with another study of our team (Li et al., 2025), our results show that no sign of aversive classical conditioning can be found in behavioral and EEG responses of individuals who are not explicitly aware of CS-US contingency. We cannot, at the present stage, generalize our findings to other kinds of associative learning, e.g., appetitive conditioning. Moreover, we cannot rule out that a different pattern of results will be observed in autonomic indices of conditioning. Because the time courses differ markedly, obtaining autonomic responses (for example, galvanic skin responses that require 4-6 s to develop) alongside much faster EEG measures, while also using many trials and maintaining a similar ratio of participants with and without contingency awareness, would be a nontrivial undertaking.

### Conclusions

Our findings indicate that associative learning in humans requires contingency awareness. Awareness plays a crucial role in anticipating and processing conditioned stimuli, as reflected in generally amplified neural patterns related to attention to informative cues, deeper sensory processing, and the violation and confirmation of expectations. Meanwhile, personality traits related to emotional processing do not influence the likelihood of becoming aware of the contingencies.

## Declarations

### Funding

The study was supported by the German Research Society (Deutsche Forschungsgemeinschaft, DFG), grant KO-1753/13-4.

### Conflicts of interest

Nothing to report.

### Ethics approval

The study was approved by the local Ethics Committee of the Faculty of Medicine of the University of Tübingen.

### Consent to participate

Participants gave their informed, written consent and were compensated with €10 or received course credit for their participation.

### Availability of data and materials

The experiment was preregistered on OSF (https://osf.io/vywq7). The data are publicly available on Openneuro (https://openneuro.org/datasets/ds005410).

### Code availability

Not available

### Author contributions statement

Yuri G. Pavlov: Conceptualization, Investigation, Data curation, Formal analysis, Project administration, Visualization, Supervision, Methodology, Writing – original draft, Writing – review & editing.

Nick S. Menger: Investigation, Writing – review & editing.

Andreas Keil: Methodology, Writing – review & editing.

Boris Kotchoubey: Conceptualization, Funding acquisition, Methodology, Supervision, Writing – review & editing.

## Supplementary

### Pilot study

Experimental design was identical to the experiment reported in the article, except the ISI (time between US presentation and start of the next trial) was 500 ms shorter (1500+/−50 ms).

#### Methods

In the pilot phase of the experiment, we collected a sample of 20 subjects (12 aware and 8 unaware).

As a result of the pilot phase of this study, we expected that about half of the participants would be able to learn the associations between the stimuli (i.e., become contingency aware).

We had several a priori hypotheses.

First, in the test phase if a first category word is presented and left-hand side vibration is expected then contralateral alpha-beta suppression preceding the actual vibration is observed. The same is applicable to the right-hand side and the second category of words.

Second, when expectation is violated e.g. the vibration is delivered to the opposite to the expected hand then error-related negativity (ERN) is generated and SPN precedes US+ presentation.

Third, the expected effects described above would be observed only in the group of aware subjects.

#### ROIs

##### ERPs

SPN1 in 500-900 ms

SPN2 in 1500-2000 ms

SPN ROI: FC1,FCz,FC2,F1,Fz,F2,C1,Cz,C2

###### Time-frequency analysis

Baseline = –0.4 –0.1

*Alpha preceding vibration*

Channels: FC3, FC1, C3, C1, CP3, CP1 (left ROI); FC2, FC4, C4, C2, CP4, CP2 (right ROI)

Time window: 600-900 ms

Frequency band: 8-14 Hz

*Beta preceding US*

Channels: FC3, FC1, FCz, FC2, FC4, C3, C1, Cz, C2, C4

Time window: 1500-2000

Frequency band: 15-22 Hz

#### Results

We did not find support for the first hypothesis. If the word associated with the left-hand side vibration was presented we did not see any lateralization of the alpha-beta suppression preceding the presentation of vibration itself. But there was a non-lateralized effect of alpha suppression in somatosensory cortex in aware group but not in unaware (see Figure S1c. The analyzed ROI is marked by black rectangle). The interaction Group x Condition was not significant, but the effect of Group was (p=0.003).

The second hypothesis was partially confirmed. We did not find anything like ERN in response to the vibrations that violate the learned rule. With that, for SPN_US_, the interaction Group (aware, unaware) x Condition (CS+ – violation of the rule and expectation of US+, CS-– vibration on the expected hand side and expectation of US-) was marginally significant (p=0.098) (see Figure S1a, circled in the right panel). SPN was more negative just before presentation of US+ comparing with expectation of US-in the aware group but not in the unaware group. The effect of Group on SPN_vibro_ (in the time window between word and vibration) presentations was also marginally significant (p=0.100) (see Figure S1a left panel).

**Figure S1.**
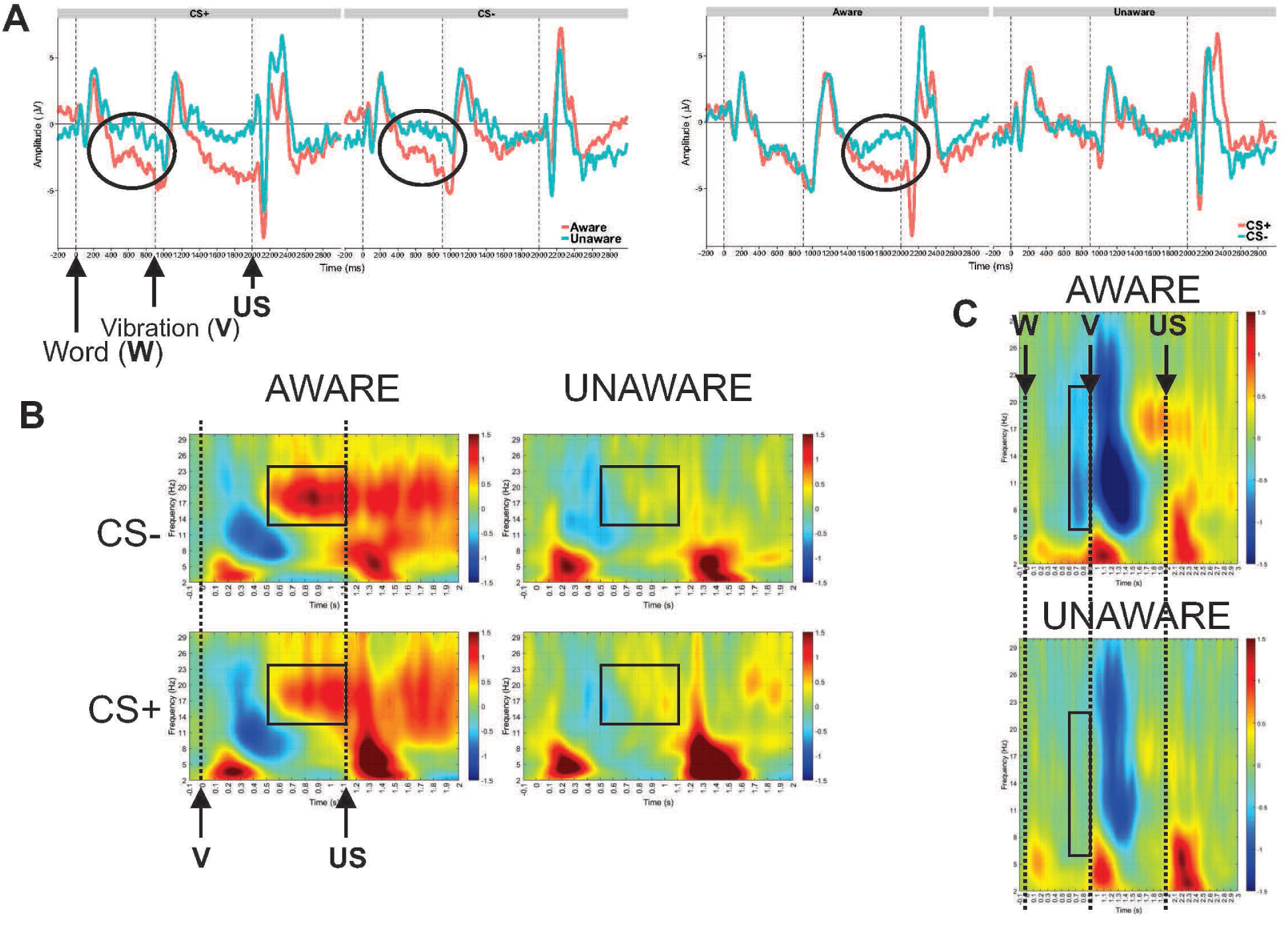
– Summary of the findings in the pilot study. (a) Left panel: Stimulus preceding negativity (SPN) at Cz is significantly different between groups (main effect of Group). Right panel: SPN is more negative in CS+ condition as compared with the CS-, but only in the Aware group (Group x Condition interaction). (b) Group effects on the oscillatory brain activity averaged over FC1, FCz, FC2, C1, Cz, C2 channels in the beta frequency band (the beta rebound effect). (c) Group effects on the alpha activity averaged over C2, C4, C6, CP2, CP4, CP6, FC2, FC4, FC6, C1, C3, C5, CP1, CP3, CP5, FC1, FC3, FC5 channels preceding the vibrations.

#### Effect sizes in the pilot study

N = 20; Group: aware (n = 12) /unaware (n=8); Condition: CS+/CS-

##### H1.1, H1.2

###### SPN2 (SPN_US_)

Channels: FC1,FCz,FC2,C2,C1,Cz,F1,Fz,F2

Time window: 1500-2000 ms after word presentation

Group *F*(1, 18) = 2.77, *p* = .114, 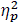 = .13

Condition *F*(1, 18) = 1.41, *p* = .251, 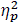 = .07

Group x Condition *F*(1, 18) = 3.05, *p* = .098, 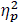 = .15

In the posthoc analysis of simple effects, the effect of condition was found to be significant only in the aware group.

Aware: t(11) = 2.320, p = 0.0323, d = 0.70

Unaware: t(7) = 0.362, p = 0.7214, d = 0.14

##### H1.3

###### SPN1 (SPN_vibro_)

Channels: FC1,FCz,FC2,C2,C1,Cz,F1,Fz,F2

Time window: 500-900 ms after word presentation

Group *F*(1, 18) = 3.01, *p* = .100, 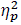 = .14

Condition *F*(1, 18) = 0.16, *p* = .692, 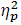 < .01

Group x Condition *F*(1, 18) = 0.12, *p* = .732, 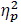 < .01

##### H1.4

###### Beta preceding US

Frequency: 15-22 Hz

Channels: FC3,FC1,FCz,FC2,FC4,C3,C1,Cz,C2,C4,F3,F1,Fz,F2,F4

Time window: 1500-2000 ms after word presentation

Group *F*(1, 18) = 4.18, *p* = .056, 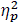 = .19

Condition *F*(1, 18) = 0.00, *p* = .978, 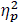 < .01

Group x Condition *F*(1, 18) = 0.47, *p* = .501, 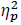 = .03

##### H1.5

###### Alpha preceding vibration

Frequency: 8-14 Hz

Channels: Pz,CPz,CP1,CP2,CP3,CP4,P1,P2,P3,P4

Time window: 500-900 ms after word presentation

Group *F*(1, 18) = 12.11, *p* = .003, 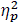 = .40

Condition *F*(1, 18) = 0.02, *p* = .880, 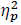 < .01

Group x Condition *F*(1, 18) = 0.10, *p* = .754, 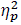 < .01

##### H1.6

###### Alpha/beta to vibration

Frequency: 8-22 Hz

Channels: Left hemisphere: C3,C1,CP1,CP3, Right hemisphere: CP2,CP4,C2,C4

Time window: 1000-1400 ms after word presentation

Group *F*(1, 18) = 4.25, *p* = .054, 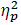 = .19

Condition *F*(1, 18) = 0.01, *p* = .907, 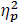 < .01

Group x Condition *F*(1, 18) = 0.03, *p* = .869, 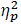 < .01

<u>Thus, the smallest Group effect was 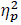 = 0.13. 75% of it is 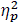 = 0.0975</u>.

#### The analysis of the acquisition phase

**Figure S2.**
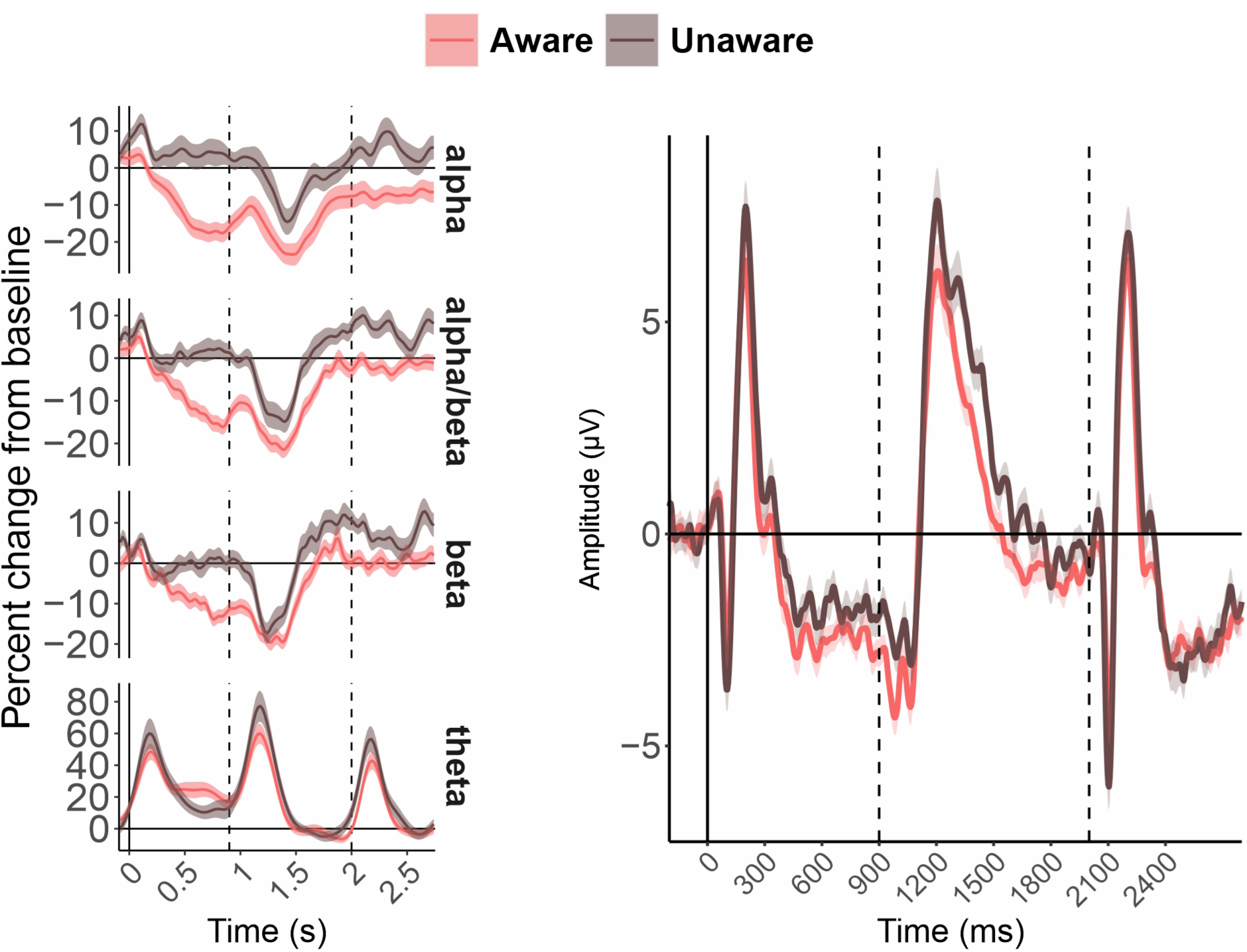
– The results of the analysis of the acquisition phase of the experiment where only CS-trials were presented. The effects in SPN_US_ and SPN_vibro_ time windows did not reach significance.

## Footnotes

1 Note that this framing is a simplification, as awareness may be graded or multidimensional, and we adopt a binary classification only for measurement clarity without claiming that the construct is inherently binary.

## Notes

https://openneuro.org/datasets/ds005410

### Competing Interest Statement

The authors have declared no competing interest.

### Summary of Updates

multiple revisions. mostly changes in the text. figures are the same

